# Supra-threshold psychoacoustics and envelope-following response relations: normal-hearing, synaptopathy and cochlear gain loss

**DOI:** 10.1101/391672

**Authors:** Sarah Verhulst, Frauke Ernst, Markus Garrett, Viacheslav Vasilkov

## Abstract

The perceptual consequences of cochlear synaptopathy are presently not well understood as a direct quantification of synaptopathy is not possible in humans. To study its role for human hearing, recent studies have instead correlated changes in basic supra-threshold psychoacoustic tasks with individual differences in subcortical EEG responses, as a proxy measure for synaptopathy. It is not clear whether the reported missing relationships between the psychoacoustic quantities and the EEG are due to the adopted methods, or to a minor role of synaptopathy for sound perception. We address this topic by studying the theoretical relationship between subcortical EEG and psychoacoustic methods for different sensorineural hearing deficits.

## 1 Introduction

The role of cochlear synaptopathy (i.e., the loss of inner-hair-cell auditory-nerve fiber synapses due to noise exposure or aging; or hidden hearing loss) for supra-threshold hearing has been heavily contested in recent human studies [1, 2, 3, 4] even though animal studies show clear histological evidence for synaptopathy [5, 6, 7]. It is not clear whether the cause of the missing correlations between subcortical EEG measures, as a non-invasive tool to quantify synaptopathy, and the suprathreshold psychoacoustic tasks stems from methodological confounds. It might be that the adopted subcortical EEG methods (e.g. the envelope-following response, EFR and auditory brainstem response, ABR) are not sensitive markers of synaptopathy in humans, or, that the EEG methods are not targeting the same mechanisms involved in the psychoacoustic task, resulting in differential effects of synaptopathy on both measures. To address these issues, we study the theoretical relationship between the EFR and two common supra-threshold hearing tasks: tone-in-noise (TiN) and amplitude-modulation (AM) detection for different degrees of sensorineural hearing loss. We employ a computational model of the human auditory periphery that simulates neural responses to quantify psychoacoustic detection cues and subcortical EEG metrics [8]. We simulate how different aspects of sensorineural hearing loss (synaptopathy, cochlear gain loss and combinations) affect the theoretical relationship between the EFR and psychoacoustic metrics to assess their sensitivity in quantifying synaptopathy in humans.

## 2 Methods

*Participants* were informed according to the ethical guidelines at Oldenburg University and paid for participation. TiN detection: 11 normal-hearing (NH; 24±4.4 yrs, 9 females) subjects with normal audiograms (Auritec AT900) and 9 hearing-impaired (HI; 63±6, 7 females) participants with a high-frequency sloping hearing loss (≤40 dB HL up to 6 kHz, with a 20 to 25 dB HL loss at 4 kHz). AM detection: 12 NH listeners (26±4, 7 females) with flat audiograms and a max. 15 dB HL threshold at 4 kHz. 8 HI listeners (70±5, 5 females) with sloping audiograms and a 4-kHz threshold between 20 and 40 dB HL.

*Psychoacoustic* stimuli were delivered monaurally using insert ER-2 earphones connected to a TDT HB7 and Fireface UCX Soundcard and were calibrated using B&K 2669, 2610, 4153, 4134 products. All measurements were conducted in a sound-proof booth and consisted of a practice run followed by a 3-alternative forced choice, 1-up-2-down procedure with 3 repetitions (AFC software). Stimuli were 500-ms long, followed by 500 ms of silence and thresholds were calculated as the mean over the last 6 reversals at the smallest step size. *TiN detection*: step sizes were 8-4-2-1 dB. A 65-dB SPL 4-kHz tone was embedded in a one-octave wide white noise masker (i.e., the reference) of varying level (SNR within one NH 4-kHz equivalent rectangular bandwidth was the tracking variable). *AM detection*: The initial modulation depth (MD) was -6 dB re 100% modulation and stepsizes were 10, 5, 3, 1 dB. The carrier was a 70-dB 4-kHz tone, the modulation frequency 100 Hz and stimulus levels for different MDs were normalized to remove loudness cues.

*EFRs* were recorded on a 32-channel Biosemi amplifier using magnetically shielded ER-2s for sound delivery while subjects watched a silent movie in a reclining chair. The 16-kHz sampled Cz data was re-referenced to the offline averaged earlobe electrodes. Each of 600 stimulus repetitions lasted 600 ms followed by a uniformly distributed random silence jitter (>90 and <110ms). Stimuli were 100% modulated 120-Hz AM signals. For the TiN experiment, EFRs were recorded to a 4-kHz centered one-octave white noise carrier of 75 dB, whereas the carrier was a 70-dB 4-kHz pure-tone for EFRs in the AM experiment. Recordings were averaged, base-line corrected and filtered between 60 and 650 Hz before epoching and bootstrapping was performed to calculate the individual noise floors and confidence intervals [9]. The FFT was calculated from the averaged -0.01 to 0.6s window re trigger onset and EFR amplitudes were calculated by adding up spectral EFR peaks (re to the noise floor) at the modulation frequency and all available harmonics (in *µ* V). The AM frequency in the psychoacoustic and EFR experiment were not identical but both greater than 80 Hz, consistent with brainstem generators of AM encoding [10].

*A computational model* of the human auditory periphery was adopted [8] to simulate a 70-dB, 4-kHz tone and 120-Hz AM tones of different modulation depths. Additionally, 20 different one-octave wide noise iterations with or without an embedded 4-kHz, 70-dB tone were averaged and simulated for a range of SNRs. Population responses were computed from simulating neural activity at the Inferior Colliculus (IC) stage of the model and by summing up time-waveforms across 401 simulated CFs spanning the human cochlear partition. Figure 1A shows an example IC population response to the 4-kHz pure tone and a 120-Hz AM tone. The *psychoacoustic detection cue* was derived from the difference signal between the IC population response to the pure-tone and AM tone. Figure 1B shows the difference signal from IC population responses to a noise and a tone embedded in noise at different SNRs. The rms of the difference signal was computed and transformed to dB to yield the detection cues plotted in Figs.2A&2B. EFRs were simulated to 100% modulated 120-Hz AM tones by adding up the population responses of the auditory-nerve (AN), cochlear nucleus and IC stages of the model to capture the different sources contributing to the scalp-recorded potential. Eight hearing profiles were simulated: (i) a NH model with normal Q_ERB_s and 3 low (1 spike/s; LS), 3 medium (10 spikes/s; MS) and 13 high (70 spikes/s, HS) spontaneous rate AN fibers synapsing at each of the 401 inner-haircells (IHCs), (ii) a selective synaptopathy model where all LS and MS fibers were removed (i.e., LS loss), (iii-iv) a synaptopathy model where all LS&MS fibers as well as 50% or 75% of the HS fibers were removed (LS50%HS and LS75%HS). Lastly, (v)-(viii) HI models with Q_ERB_s corresponding to a high-frequency sloping audiogram (above 1-kHz) up to 35 dB at 8 kHz and synaptopathy profiles as in (i)-(iv).

**Figure 1:**
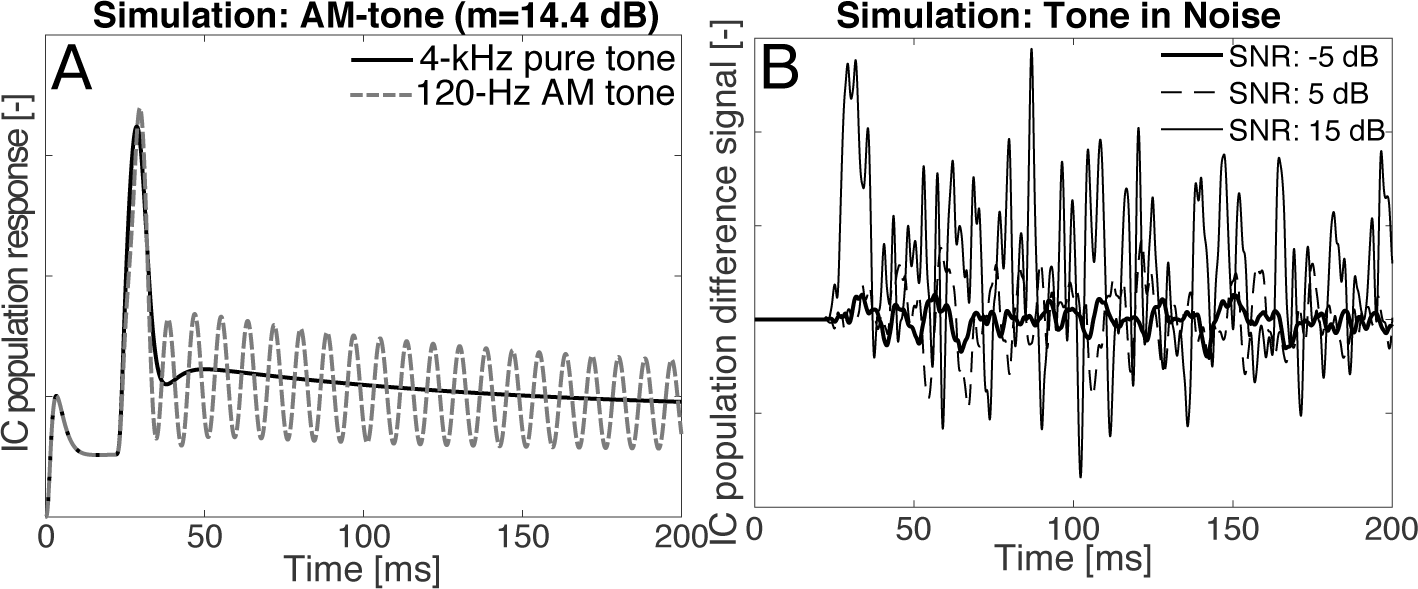
**A**: Simulated IC population responses to a 4-kHz pure tone and a 120-Hz, 4-kHz AM tone of 70 dB SPL. The detection cue is the rms of the difference signal of the waveforms. **B**: Difference signal between simulated responses for noise and TiN epochs of a 70dB-SPL 4-kHz pure tones embedded in a one-octave wide white noise for different SNRs. The rms of the difference signal is the simulated detection cue.

## 3 Results

### Psychoacoustics

Figures 2A&B depict simulated psychoacoustic detection cues for the AM and TiN detection experiment and show that synaptopathy has a greater influence on shifting the NH curve downward than a high-frequency sloping cochlear gain loss. In fact, the AM detection cue is somewhat stronger in the HI models for the same degree of synaptopathy. In the model, this is explained by a lower effective drive to the IHC-AN complex caused by cochlear gain loss, resulting in less saturated AN responses and enhanced AM sensitivity, corroborating observations in the chinchilla AN [11]. The psychoacoustic threshold for the NH model was set to the detection cue corresponding to the modulation depth at which NH people performed (i.e., -29.5 dB; 2^*nd*^ best human NH performer in Fig.3A). The gray threshold line in Fig. 2A shows that the AM threshold shifted by 8 dB and even by 15 dB for the LS50%HS and LS75%HS models respectively. Similarly, Fig.2B predicts the need for a 4-dB stimulus SNR increase for the LS50%HS models to reach the reference NH detection cue amount and performance.

**Figure 2:**
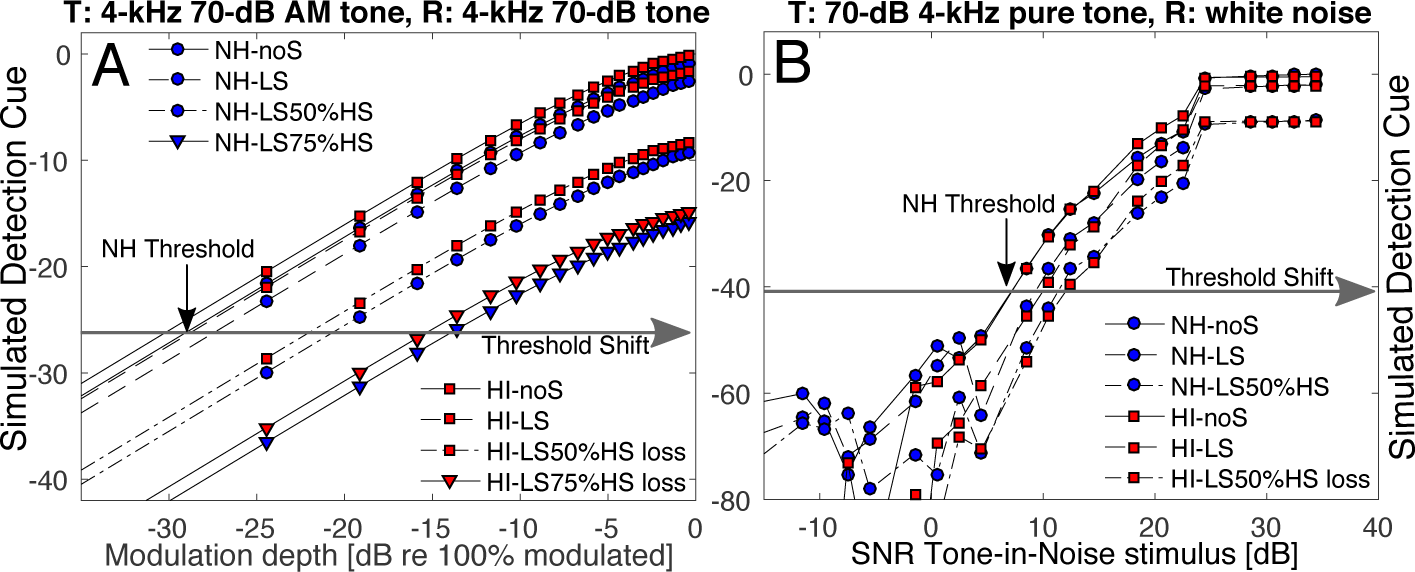
(Colour online) **A**: Simulated AM detection cues for different hearing loss models. LS: selective LS&MS fiber loss, LS50%HS: LS&MS and 7 HS fibers lost at each CF, LS75%HS: LS&MS and 10 HS fibers lost at each CF. The threshold line shows the shift from the reference NH AM detection threshold (-29.5 dB) **B**: Simulated TiN detection cues. The threshold line indicates the SNR at the detection threshold shift in dB from the NH human reference SNR (7.2 dB).

Figures 3A&B summarize the simulated detection cue shifts (black lines, filled markers) and EFR amplitude reductions (colored lines, filled markers) alongside human reference data (open markers) for NH and HI participants who performed significantly worse on all measures (p<0.01). As the simulated detection cue shifts for the HI models were similar to those of the NH models, we conclude that synaptopathy rather than cochlear gain loss was responsible for the degraded detection cues. The range of simulated AM detection thresholds caused by synaptopathy and the spread in the reference data corresponded well. The best thresholds in the reference AM experiment were in line with those in [12] and the data-spread was about 15-20 dB across listeners where NH aging studies predict age-related reductions between 5 and 10 dB in the absence of cochlear gain loss [13, 14]. Our simulations suggest that synaptopathy can explain a large degree of the individual performance despite co-existing elevated hearing thresholds. The absence of a relationship between the experimental 4-kHz pure-tone and AM detection thresholds supports this notion (R^2^=0.3; p=0.09). The *≈*7dB spread in the TiN reference data corresponded well to the shift predicted by synaptopathy and corroborate reported 5-10 dB TiN threshold shift in a fixed 50-dB-SPL broadband noise when more than 60% of the IHC population is lost [15]. However, in contrast to the AM thresholds, degraded TiN detection performance was also related to elevated pure-tone thresholds (R^2^=0.3; p=0.02).

**Figure 3:**
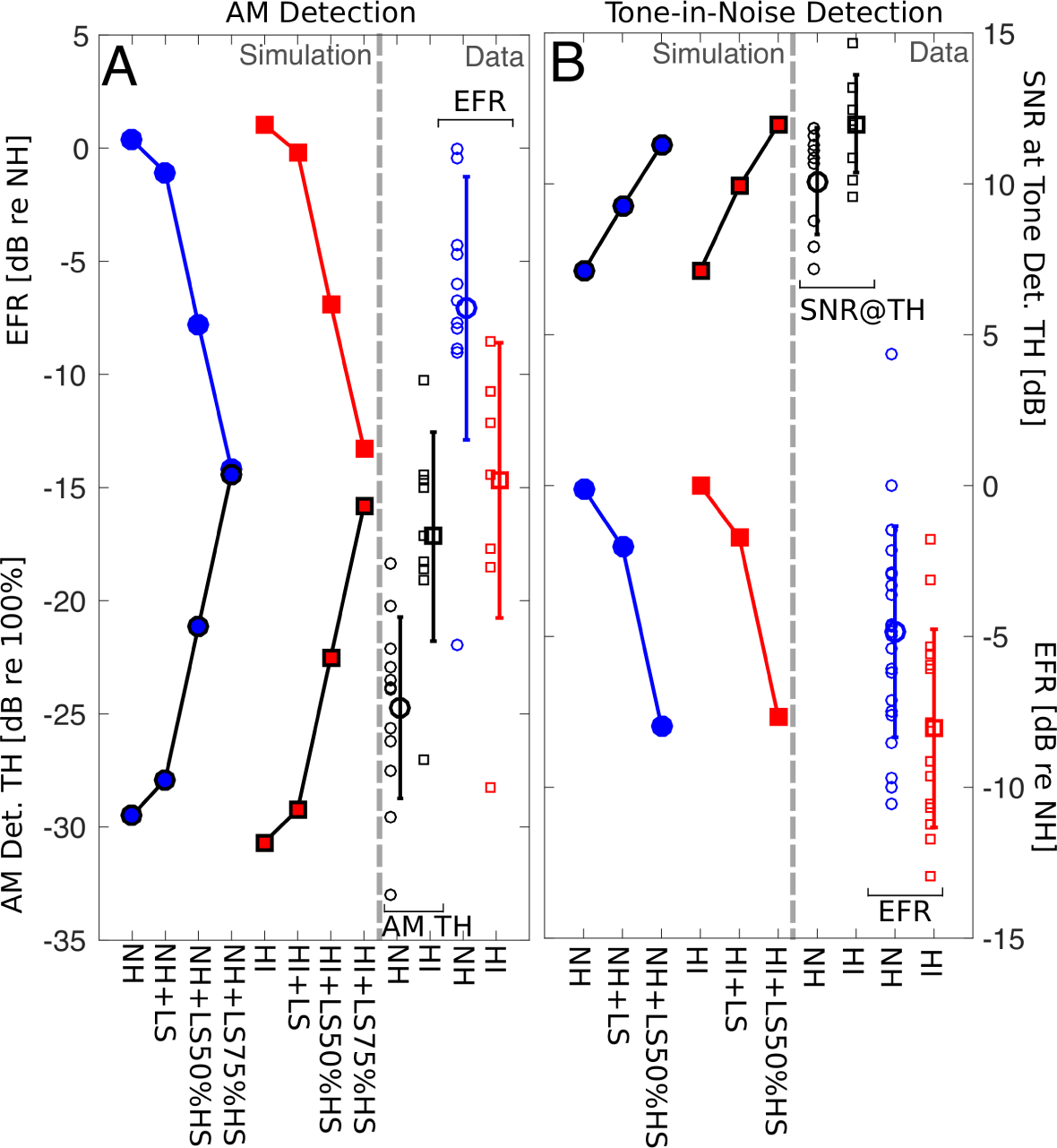
(Colour online) **A**: Simulated and recorded AM detection thresholds (70-dB, 4-kHz AM tone) and EFRs to 100% modulated 120-Hz 4-kHz tones [in dB re best NH EFR] for different hearing loss profiles. **B**: Simulated tone-in-noise detection thresholds (70-dB SPL tone) and EFRs to 100% modulated 120-Hz 4-kHz tones [in dB re best NH EFR] for different hearing-loss profiles. Reference EFRs were recorded to 75-dB SPL, one-octave wide, 4-kHz centered 100%, 120-Hz modulated white noises.

### Relation between psychoacoustics and EFRs

Figure 3 compares simulated and recorded EFR amplitudes re the reference NH EFR amplitude (blue and red). The spread of simulated and recorded EFR amplitude reductions around their mean coincide, although the HI EFRs showed overall greater reductions than predicted, suggesting that synaptopathy differences may only partly explain the individual spread in the human data. To test whether psychoacoustic metrics can be used as a replacement for EFRs in the diagnosis of synaptopathy, we studied their relationship. While AM detection and EFRs both rely on a robust coding of temporal envelope information, the sensitivity to small modulation depths (the psychoacoustic task) may not be a predictor of the EFR amplitude to 100% modulated stimuli. Regression fits between individual EFR and psychoacoustic metrics (irrespective of their NH or HI status) are depicted in Fig. 4. The best simulated NH psychoacoustic threshold was matched to that of the best performing NH listener, while simulated EFR amplitudes were not scaled in the analysis. Simulated detection cues and EFR amplitudes related well (R^2^ >0.9) and the fit was somewhat better for the AM detection task due to stimulus similarity. In the model, the regression is generally predictive of the degree of synaptopathy (not cochlear gain loss). The experimental results show a larger spread around the regression line (R^2^ of 0.3 and 0.4) than predicted by the model, but nevertheless show a significant relationship. The regression for the AM experiment extend the reported NH correlation between the EFR and AM detection [1] to HI listeners. The experimental relationship between tone-in-noise detection SNR at threshold and the EFR amplitude has not been reported earlier, but its existence is supported by the model simulations.

**Figure 4:**
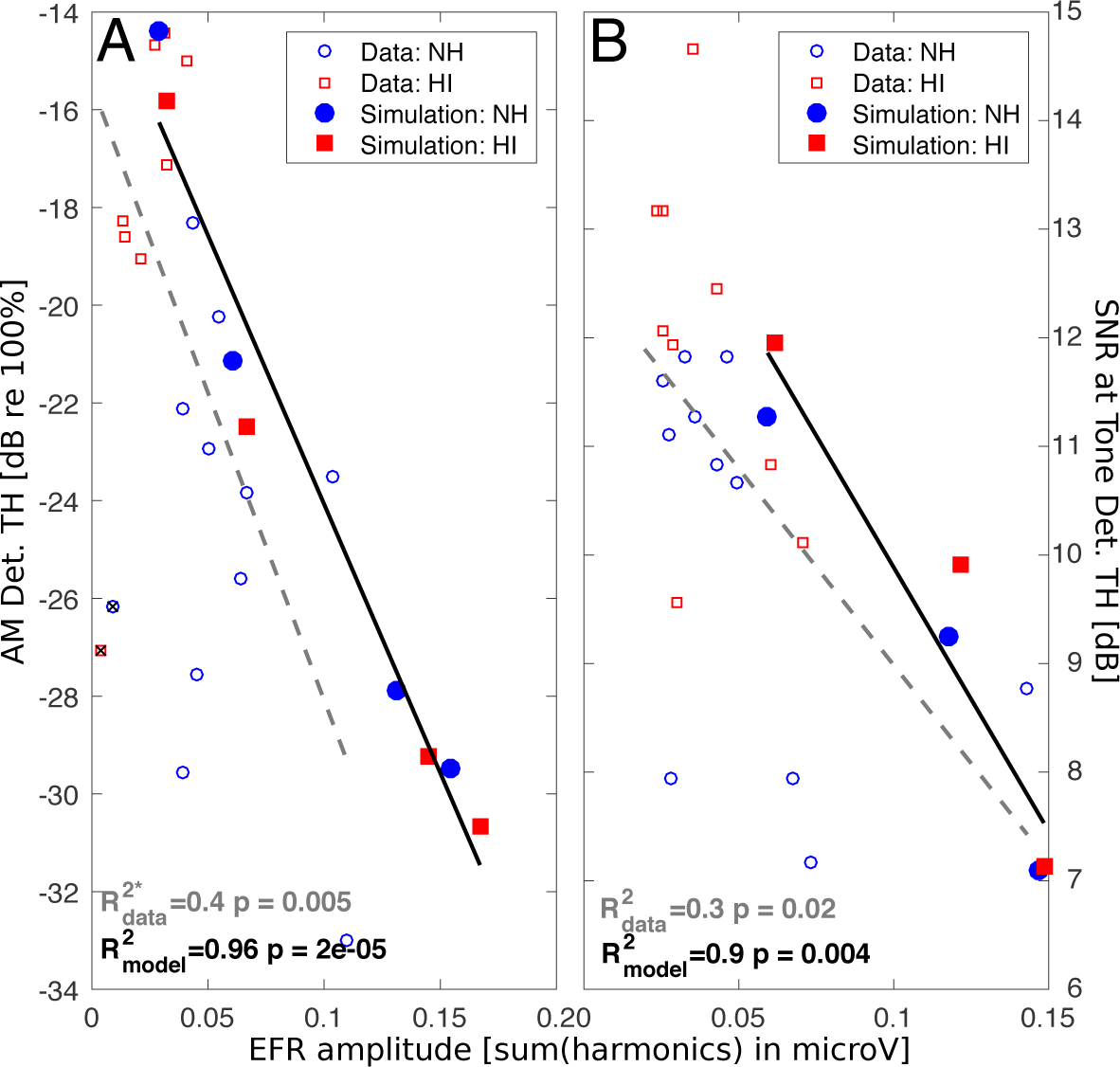
(Colour online) Regression for simulated and recorded NH and HI measures. Model simulations (filled circles and squares) refer to no-synaptopathy (bottom marker), LS, LS50%HS & LS75%HS (top marker). **A**: AM detection threshold for 70 dB AM (100/120Hz) tone. The two crossed symbols were excluded from the regression analysis as the EFR amplitudes (shown) were much less strong than those derived for the −4 or −8 dB conditions (not shown), pointing to a problem with the EFR recordings. **B**: SNR at tone-detection threshold for a 65/70-dB, 4-kHz pure tone in white noise.

## 4 Discussion

Both supra-threshold psychoacoustic tasks were strongly affected by synaptopathy and only mildly by cochlear gain loss for the considered stimulus configurations, suggesting that these tasks may differentially diagnose synaptopathy in NH *and* HI listeners. Even though simulations are inherently limited by the quality of the model (which does not account for plasticity or cognitive factors), we propose that the effect of synaptopathy on supra-threshold psychoacoustsics is much greater than so far assumed. Signal detection theory predicts a 1.5-dB or 5-dB shift in the TiN detection threshold for a respective loss of 50% and 90% of the available AN fibers [2], whereas we observed that a 70% fiber loss (i.e., LS50%HS) in a functional model of the human auditory 1 periphery causes a threshold shift of 4 dB. For AM detection, a 70% or 85% (i.e., LS75%HS) fiber loss predicted 1 a respective 8 and 15-dB threshold shift, which matched the individual variability in the combined NH and HI 1 reference data well. Controversially, we propose that the reason why the HI listeners performed worse than the NH listeners, was due to their synaptopathy and AN fiber loss and not because of their coexisting outer haircell loss deficits. This latter aspect can be confirmed experimentally, as age-related synaptopathy was shown to occur before outer haircell loss [16]. If our predictions are correct, ageing listeners with normal audiometric thresholds suffering from synaptopathy should show EFRs, TiN and AM detection thresholds in range with those of HI participnts.

## Acknowledgements

European Research Council grant agreement No 678120 (SV,VV). DFG Cluster of Excellence EXC 1077-1 “Hearing4all” (MG) and DFG PP 1608 VE924/1-1 (FE).

## References

[1] H.M. Bharadwaj, Masud, Mehraei, Verhulst, Shinn-Cunningham: Individual differences reveal correlates of hidden hearing deficits. J Neurosci 35(5) (2015) 2161–2172.

[2] A.J. Oxenham: Predicting the perceptual consequences of hidden hearing loss. Trends Hear. 20 (2016) 2331216516686768.

[3] G. Prendergast, Millman, Guest, Munro, Kluk, Dewey, Hall, Heinz, Plack: Effects of noise exposure on young adults with normal audiograms II: Behavioral measures. Hear Res 356 (2017) 74–86.

[4] I. Yeend, Beach, Sharma, Dillon: The effects of noise exposure and musical training on suprathreshold auditory processing and speech perception in noise. Hear Res 353 (2017) 224–236.

[5] S.G. Kujawa, Liberman: Adding insult to injury: cochlear nerve degeneration after “temporary” noise-induced hearing loss. J Neurosci 29(5) (2009) 14077–14085.

[6] D. Möhrle, Ni, Varakina, Bing, Lee, Zimmermann, Knipper, Rüttiger: Loss of auditory sensitivity from inner hair cell synaptopathy can be centrally compensated in the young but not old brain. Neurobiol Ageing 44 (2016) 173–184.

[7] M.D. Valero, Burton, Hauser, Hackett, Ramachandran, Liberman: Noise-induced cochlear synaptopathy in rhesus monkeys (Macaca mulatta). Hear Res 353 (2017) 213–223.

[8] S. Verhulst, Altoè, Vasilkov: Computational modeling of the human auditory periphery: Auditory-nerve responses, evoked potentials and hearing loss. Hear Res 360 (2018) 55–75.

[9] M. Garrett, Verhulst: Applicability of cochlear synaptopathy sensitive subcortical EEG metrics in the normal and impaired auditory system. Trends Hear (in review).

[10] D. Purcell, John, Schneider, Bruce, Picton: Human temporal auditory acuity as assessed by envelope following responses. J Acoust Soc Am 116(6) (2004) 3581–3593.

[11] S. Kale, Heinz: Envelope coding in auditory nerve fibers following noise-induced hearing loss. J Assoc Res Otolaryng 11(6) (2010) 657–673.

[12] A. Kohlrausch, Fassel, Dau: The influence of carrier level and frequency on modulation and beat-detection thresholds for sinusoidal carriers. J Acoust Soc Am 108(2) (2000) 723–734.

[13] S. Jin, Liu, Sladen: The effects of aging on speech perception in noise: comparison between normal-hearing and cochlear-implant listeners. J Am Ac Audiol 25(7) (2014) 656–665.

[14] G. Takahashi, Bacon: Modulation detection, modulation masking, and speech understanding in noise in the elderly. J Speech Language Hear Res 35(6) (1992) 1410–1421.

[15] E. Lobarinas, Salvi, Ding: Selective inner hair cell dysfunction in chinchillas impairs hearing-in-noise in the absence of outer hair cell loss. J. Assoc. Res. Otolaryngol 17(2) (2015) 89–101.

[16] Y. Sergeyenko, Lall, Liberman, Kujawa: Age-related cochlear synaptopathy: an early-onset contributor to auditory functional decline. J Neurosci 33(34) (2013) 13686–13694.

